# Encoding of interdependent features of head direction and angular head velocity in navigation

**DOI:** 10.1101/2024.05.10.593505

**Authors:** Dongqin Cai, Tao Liu, Jia Liu

## Abstract

To comprehend the complex world around us, our brains are tasked with the remarkable job of integrating multiple features into a cohesive whole. While previous studies have primarily focused on the processing and integration of independent features, here we investigated the simultaneous encoding of the interdependent features, specifically head direction (HD) and its temporal derivative, angular head velocity (AHV), by first employing computational modeling on HD systems to explore emergent algorithms and then validated its biological plausibility with empirical data from mice’s HD systems. Our analysis revealed two distinct neuron populations: those with multiphasic tuning curves for HD compromised their HD encoding capacity to better capture AHV dynamics, while those with monophasic tuning curves primarily encoded HD. This pattern of functional dissociation was observed in both artificial HD systems and the cortical and subcortical regions upstream of biological HD systems, suggesting a general principle for encoding interdependent features. Further, exploration of the underlying mechanisms involved examining neural manifolds embedded within the representational space constructed by these neurons. We found that the manifold by neurons with multiphasic tuning curves was locally jagged and complex, which effectively expanded the dimensionality of the neural representation space and in turn facilitated a high-precision representation of AHV. Therefore, the encoding strategy for HD and AHV likely integrates characteristics of both dense and sparse coding schemes to achieve a balance between preserving specificity for individual features and maintaining their interdependency nature, marking a significant departure from the encoding of independent features and thus advocating future research delving into the encoding strategies of interdependent features.

## INTRODUCTION

Consider the simple act of bicycling along a winding path in a park. As one navigates a curve, the brain must simultaneously process the direction in which the head is turning (head direction, HD) and the speed of this turn (angular head velocity, AHV) to ensure a smooth ride despite the path’s twists and turns. Previous studies have primarily focused on the processing and integration of independent features (Campo et al., 2021; Spence, 2020), such as smell, touch, taste, and sight, to fabricate the unified experience of enjoying a cup of coffee. This integration typically involves neurons that exhibit either pure selectivity, allowing for precise processing and interpretation of specific features (Vaccari et al., 2022; Weinberger, 1995), or mixed selectivity, encoding combinations of multiple features to enhance the brain’s computational flexibility and efficiency (Fusi et al., 2016; Kira et al., 2023; Ledergerber et al., 2021; Ma et al., 2023; Rigotti et al., 2013). Nonetheless, the simultaneous encoding of interdependent features, exemplified by HD and its temporal derivative, AHV, poses a great challenge, which requires a delicate balance between preserving specificity for each individual feature to prevent mutual interference and maintaining their interdependency nature, given the crucial role of AHV in updating HD. Here, we asked how the brain encodes these interdependent features simultaneously.

Previous studies have identified numerous cortical regions across mammals that are involved in the encoding of HD and AHV. HD neurons are predominantly located within the limbic (Cho and Sharp, 2001; Giocomo et al., 2014; Stackman and Taube, 1998; Taube, 1995; Taube et al., 1990a, 1990b) and extralimbic areas (Hennestad et al., 2021; Keshavarzi et al., 2022; Long et al., 2022; Mehlman et al., 2019), whereas AHV neurons are identified in early peripheral nuclei such as the lateral mammillary nucleus (Stackman and Taube, 1998) and dorsal tegmental nucleus (Bassett and Taube, 2001). Recent studies reveal that AHV neurons are co-distributed with HD neurons in the motor (Mehlman et al., 2019), visual (Vélez-Fort et al., 2018), and retrosplenial cortices (Keshavarzi et al., 2022), demonstrating a brain-wide conjunctive encoding of HD and AHV (Brennan et al., 2021; Hennestad et al., 2021; Keshavarzi et al., 2022; Taube, 1995).

Inspired by the coding scheme for multiple independent features, we proposed two related but distinct theoretical hypotheses regarding the conjunctive encoding of HD and AHV (Fig. 1). The first hypothesis suggests a coding scheme analogous to neurons with mixed selectivity, referred to as “dense coding”. In this scheme, a single neuron responds to all features (e.g. (Finkelstein et al., 2018; Knierim and Zhang, 2012; Vafidis et al., 2022)), revealing the interdependency between HD and AHV at both neuronal and population levels (Fig. 1A). The second hypothesis proposes a coding scheme similar to neurons with pure selectivity, referred to as “sparse coding”. In this scheme, multiple heterogeneous neurons exhibit varying preferences for either HD or AHV (Jacob et al., 2017; Kornienko et al., 2018; Olson et al., 2017), and the interdependency between HD and AHV is only achieved through the orchestrated interaction of these heterogeneous neurons (Fig. 1B). Here, we test these two coding schemes by first using computational modeling on HD systems to explore algorithms that simultaneously encode the interdependent variables of HD and AHV, and then analyzing empirical data from mice’s HD systems to verify the biological plausibility of these algorithms.

**Figure 1.**
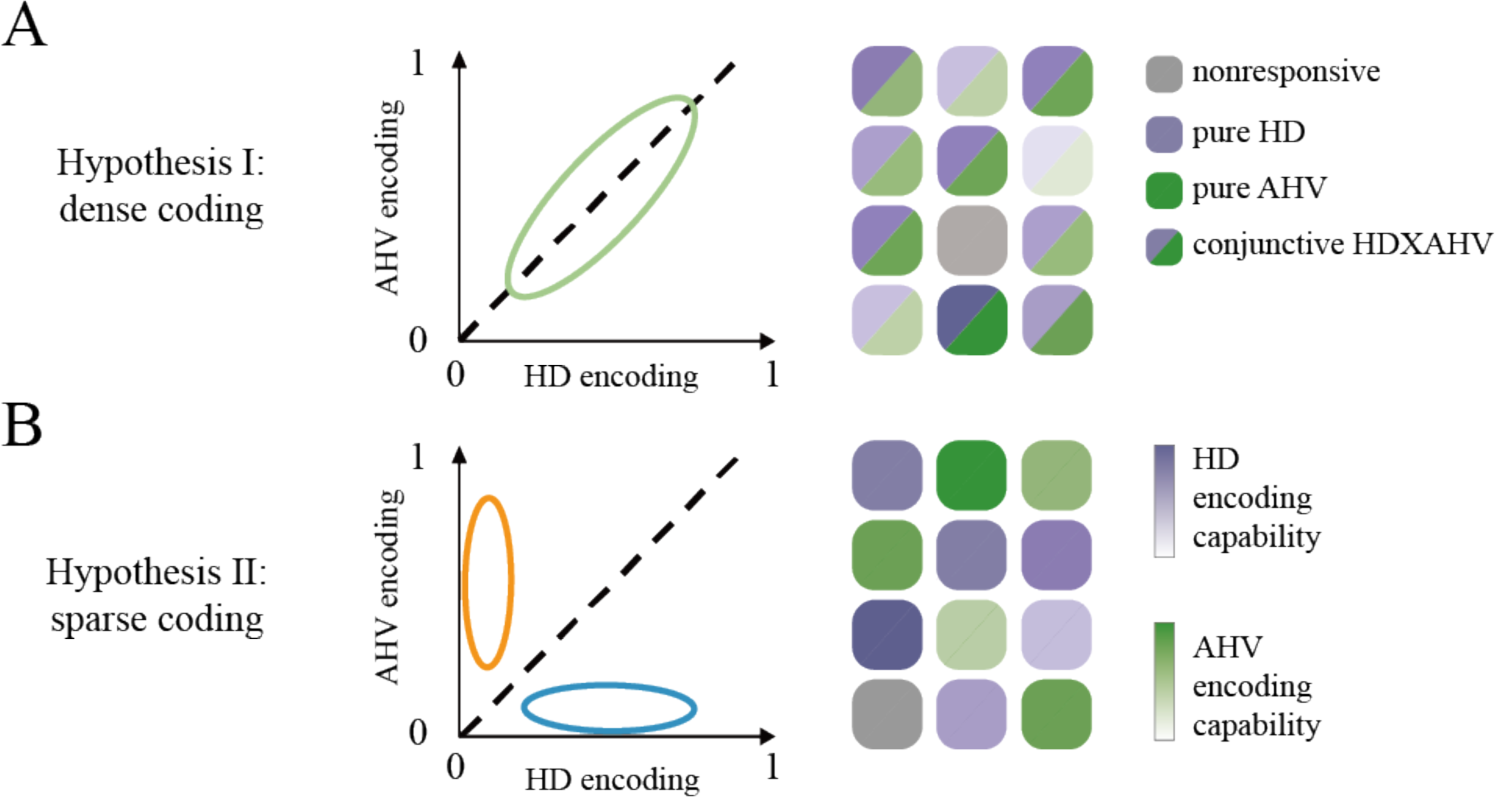
Two hypothetic coding schemes for simultaneously encoding HD and AHV. (A) Dense coding. HD and AHV are encoded by a population of homogeneous neurons that exhibit varying sensitivities to both variables, with the interdependency between HD and AHV evident both at the neuronal level, through mixed selectivity response profiles, and at the population level, through discernible activation patterns. (B) Sparse coding. HD and AHV are encoded by two distinct neuron types, each specializing in either HD or AHV. The interdependency is primarily observed at the population level, where the collective activity patterns of these distinct neurons reveal the integrated representation of both variables. The gradient from light to dark colors alongside the 0-1 range on both axes visually represents the spectrum of encoding efficacy within this neuronal population, with darker colors and values closer to 1 indicating superior encoding performance. Each square denotes the response profile of an individual neuron.

Specifically, we first explored self-emergent algorithms through an encapsulated computational model specifically designed to achieve the computational goal of simultaneously encoding HD and AHV, where similar models in previous studies have revealed the conjunctive coding of task-related variables (Cueva et al., 2019; Low et al., 2023). This approach mitigates potential confounding effects from interactive influences of upstream and downstream cortical regions or from intrinsic functionalities not directly related to HD and AHV within biological HD systems. Our modeling revealed two types of units with monophasic and multiphasic tuning curves, specialized in encoding HD and AHV, respectively. This pattern suggests a combination of the two coding schemes that balance the specificity and the interdependency between HD and AHV. Then, we validated this coding algorithm with empirical data from mice’s cortical and subcortical regions, confirming the functional division of labor in encoding HD and AHV among these neuron types, suggesting a general principle within HD systems. Finally, we investigated the underlying mechanisms by analyzing neural manifolds embedded in the neural representational space constructed by these two neuron types, revealing that neurons with multiphasic tuning curves effectively expand the dimensionality of the neural representation space, facilitating high-precision representations of AHV, essentially the temporal derivative of HD.

## RESULTS

### Emergence of single-peaked and multiple-peaked units for HD in RNN of HD system

Here we employed a recurrent neural network (RNN) model (Fig. 2A, top), which mirrors the anatomical and functional characteristics of neurons within the HD system observed in rodents and fruit flies (Cueva et al., 2019), to uncover the self-emergent algorithms that allow for the simultaneous encoding of these two interdependent variables. Besides its recurrent architecture that models neural interconnections, the dynamics of each unit in the RNN are regulated by the firing rate model (Bosio et al., 1991), simulating the firing characteristics of neurons. The model input consists of three terms: two inputs encoding the initial HD in the form of sin θ_0_ and cos θ_0_, and a scalar input encoding either clockwise (negative) or counterclockwise (positive) AHV at every timestep. The output of the model is the predicted HD based on the input. The model was trained using the Hessian-free algorithm (Martens and Sutskever, 2011) and optimized through stochastic gradient descent to minimize the mean-squared error between the predicted HDs from the RNN model and the actual HDs derived directly through integration of AHV.

**Figure 2.**
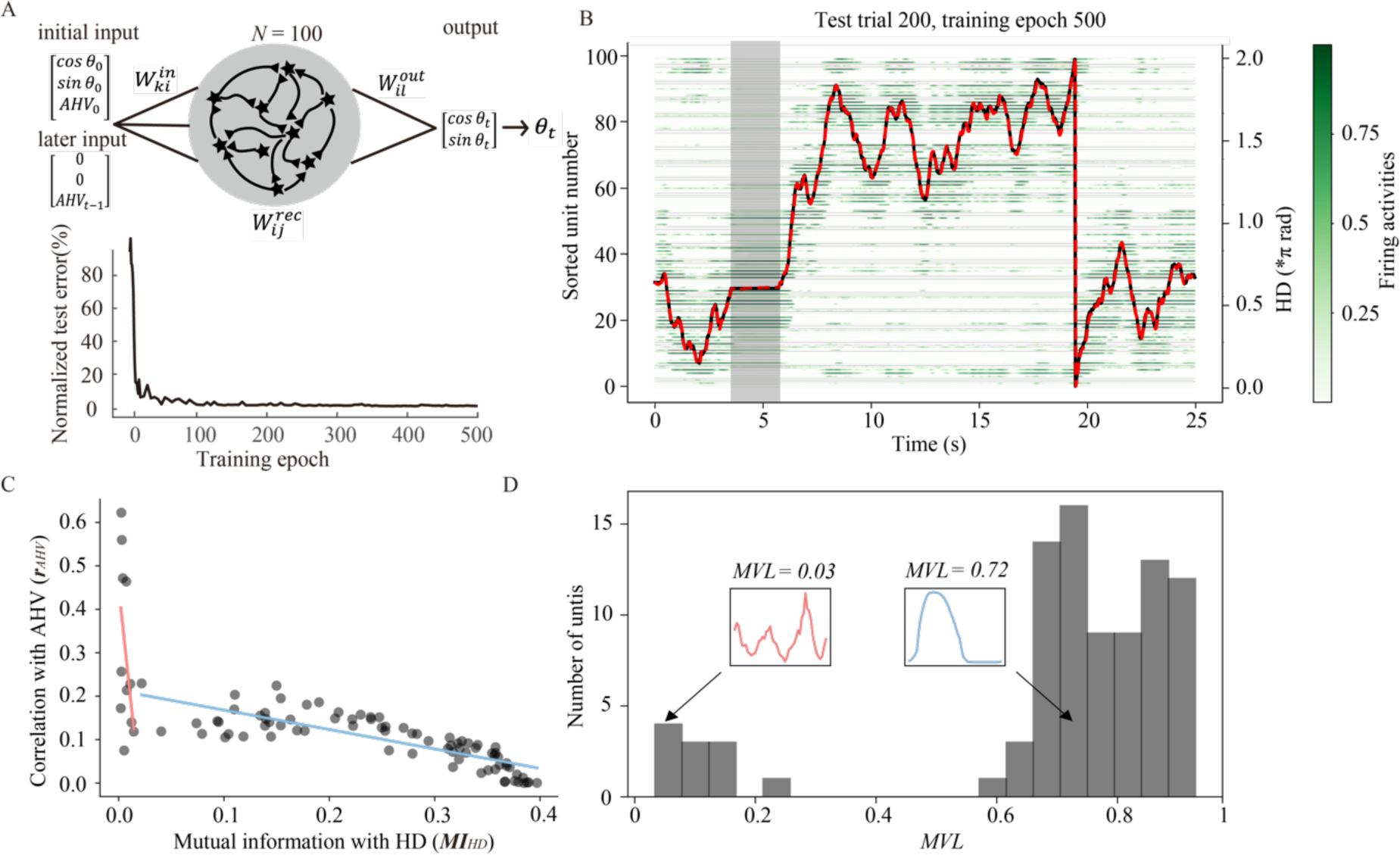
RNN model and the emergence of single-peaked (SP) and multiple-peaked (MP) units. (A) Top: a schematic diagram illustrating the RNN model trained to predict real-time HD θ_&_using initial HD and real-time AHV as inputs. Bottom: Mean-squared errors on testing trials over training epochs normalized by that of the first epoch. (B) Superimposed traces of predicted HD (red), ground truth HD (black), and firing rates (green) for all units in an example test trial. The shaded region shows the predicted HD maintained stable when AHV was set to 0. Y axis denotes individual units, with the intensity of green color indicating normalized firing rates. (C) Relationship between the encoding strengths of HD and AHV for each individual unit, showing two distinct populations of units. Linear regression lines are shown for each population. Each dot denotes an individual unit. (D) Histogram of *MVL* values for all units significantly tuned to HD. Insets display exemplar SP and MP tuning curves with corresponding *MVL* values.

For each training epoch, the input AHV was from simulated datasets with distributions matched to those observed in rats (Sharp et al., 2001; Stackman and Taube, 1998). In total, 500 trials of 500 timesteps were used for training, and the updated model parameters were tested on another 500 new trials of 500 timesteps to calculate the mean-squared error between the predicted and actual HDs. The errors in the test trials decreased drastically in the first 10 epochs and reached near the minimum around 300^th^ epoch (Fig. 2A, bottom). After extensive training for 500 epochs, the RNN model showed exceptional performance in tracking changes in HD in response to varying AHV inputs (Fig. 2B). In addition, the model also maintained a stable HD representation when AHV was set to 0 (Fig. 2B, the shaded region). We thus used the model to generate 20,0000 timestep samples as one simulated session for subsequent analyses.

We examined how HD and AHV were encoded at both single-unit and population levels with all units included, an advantage afforded by computational modeling. We assessed a unit’s capacity of encoding HD by calculating the mutual information (Skaggs et al., 1993) between HD and averaged firing rates, referred to as *MI_HD_*. This metric stands for shared entropy between two dependent variables (Hennestad et al., 2021; Peyrache et al., 2017), with a larger value indicating better capacity of encoding HD. Meanwhile, a unit’s capacity of encoding AHV was characterized by *r_AHV_*, which is the absolute values of Pearson’s correlation coefficients between the AHV and the averaged firing rates (Keshavarzi et al., 2022; Long et al., 2022), where positive and negative velocities were considered separately. A higher value indicates better capacity of encoding AHV. We found that a significant proportion of units (84%) in the RNN exhibited varying degrees of sensitivity to both HD and AHV (Fig. 2C), replicating the finding of diverse encoding capabilities across neurons in previous studies (Cueva et al., 2019; Hennestad et al., 2021). Specifically, there was a significant negative correlation between *MI_HD_* and *r_AHV_* (r = -0.67, *p* < 0.001), which suggested that units adept at encoding HD tended to have poor performance in encoding AHV. Upon closer examination, we found that the decline in *r_AHV_* was more pronounced at the lower levels of *MI_HD_* compared to the higher levels (Fig. 2C). This pattern was accurately modelled by a two-stage piecewise linear function, with a steeper fitted slope (*ρ* = -16.9) for *MI_HD_* values below 0.025, and a gentler slope (*ρ* = -0.28) for *MI_HD_* values above 0.025. This model demonstrated a superior fit (*R^2^*= 0.62) compared to other models that did not account for bifurcation (Quadratic, *R^2^* = 0.47; Exponential, *R^2^* = 0.48; Linear, *R^2^* = 0.45), suggesting the presence of two distinct populations of units with different conjunctive coding patterns for HD and AHV.

Based on the finding that a neuron’s tuning curve pattern critically influence its information encoding capacity (Kriegeskorte and Wei, 2021; Lenninger et al., 2023), here we examined whether the two distinct populations demonstrated unique tuning characteristics. To do this, all units’ tuning curves for HD were constructed based on their firing rates across all timesteps (Supplementary Fig. 1A and 1B). We found a notable divergence in *MI_HD_* values associated with the tuning curve patterns: units with higher *MI_HD_* predominantly exhibited monophasic tuning curves, marked by a concentrated distribution of HDs that elicited elevated firing rates (units with blue tuning curves in Supplementary Fig. 1A). In contrast, units with lower *MI_HD_* were found to exhibit multiphasic response profiles, displaying multiple, distinct peaks corresponding to varied HDs (units with red tuning curves in Supplementary Fig. 1A). To quantify this observation, we calculated the mean vector length (*MVL* or Rayleigh r) for each unit (Long et al., 2022; Yoder and Taube, 2009), with values ranging from 0 and 1. Larger *MVL* value signifies a unit’s firing concentration around a particular HD, indicating a monophasic selectivity, whereas lower *MVL* value suggests a unit’s mixed selectivity for multiple HDs, indicative of a multiphasic response profiles. The distribution of *MVL* values across all units showed a bimodal pattern (Fig. 2D), with a dominant portion (87.5%) of units demonstrating monophasic tuning curves (*MVL* > 0.5), hence labeled as single-peaked (SP) units. In contrast, units displaying multiphasic tuning curves (*MVL* < 0.5) constituted about 10% of the population, labeled as multiple-peaked (MP) units. This small but non-negligible proportion of MP units persisted even after downsizing the RNN model (Supplementary Fig. 1D and E), highlighting a consistent structural dichotomy. Following this observation, we further explored the intricate relationship between the structural attributes of tuning curves and the encoded HD and AHV they conveyed.

To quantify the encoding capabilities for HD and AHV of these two distinct unit types, we employed a decoding approach utilizing deep feedforward artificial neural network (ANN) (for details, see Methods), known for their effectiveness in neural decoding tasks by leveraging higher-order nonlinearities and offering end-to-end training to maximize information extracted from neural firing patterns (Ajabi et al., 2023; Richards et al., 2019; Xu et al., 2019). For each simulated RNN session, one ANN was trained to decode either HD or AHV values from the neural activity of either SP or MP unit population. The Pearson’s correlation coefficient, referred as decoding *r*, between the ANN-predicted and actual HD or AHV values, served as a metric for the population’s encoding capability. Based on testing results on our own dataset and previous studies (Ito et al., 2022; Xu et al., 2019), this decoding method demonstrated high reliability, exhibiting strong generalization from training to testing datasets and minimal variance across different cross-validation splits and simulation sessions.

Fig. 3A and Fig. 3B show the correlation between the predicted and actual data during a representative simulated session revealed distinct encoding capacities for HD and AHV, respectively, between the SP and MP unit populations. The SP unit population demonstrated superior encoding for HD than the MP unit population (decoding *r*: 0.99 versus 0.96), consistent with previous studies associating lower decoding accuracy with neurons characterized by more uniformly distributed tuning curves (Xu et al., 2019). In contrast, the MP unit population exhibited an enhanced capacity for AHV representation (decoding *r*: 0.90 versus 0.81). This dissociation was further illustrated through the analysis of decoding accuracy distribution across 50 simulated sessions with random seeds (Fig. 3C), showing an opposite pattern of encoding capacities between SP and MP unit populations (SP versus MP in HD: 1.00 versus 0.96, *p* < 0.001, Mann-Whitney U Test; SP versus MP in AHV: 0.81 versus 0.90, *p* < 0.001, Mann-Whitney U Test), which was further confirmed by a two-way interaction of unit population (SP versus MP units) by encoded variable (HD versus AHV) (F(1, 98) = 17572, *p* < 0.001). Another way to segregate units into distinct populations is based on varying *MVL* thresholds. We found that even a minimal cohort of MP units with the lowest *MVL* values outperformed the remainder possessing higher *MVL* values in encoding AHV. In contrast, a population of SP units with the highest *MVL* values approached a decoding *r* near 1, indicative of near-perfect HD encoding (Fig. 3D).

**Figure 3.**
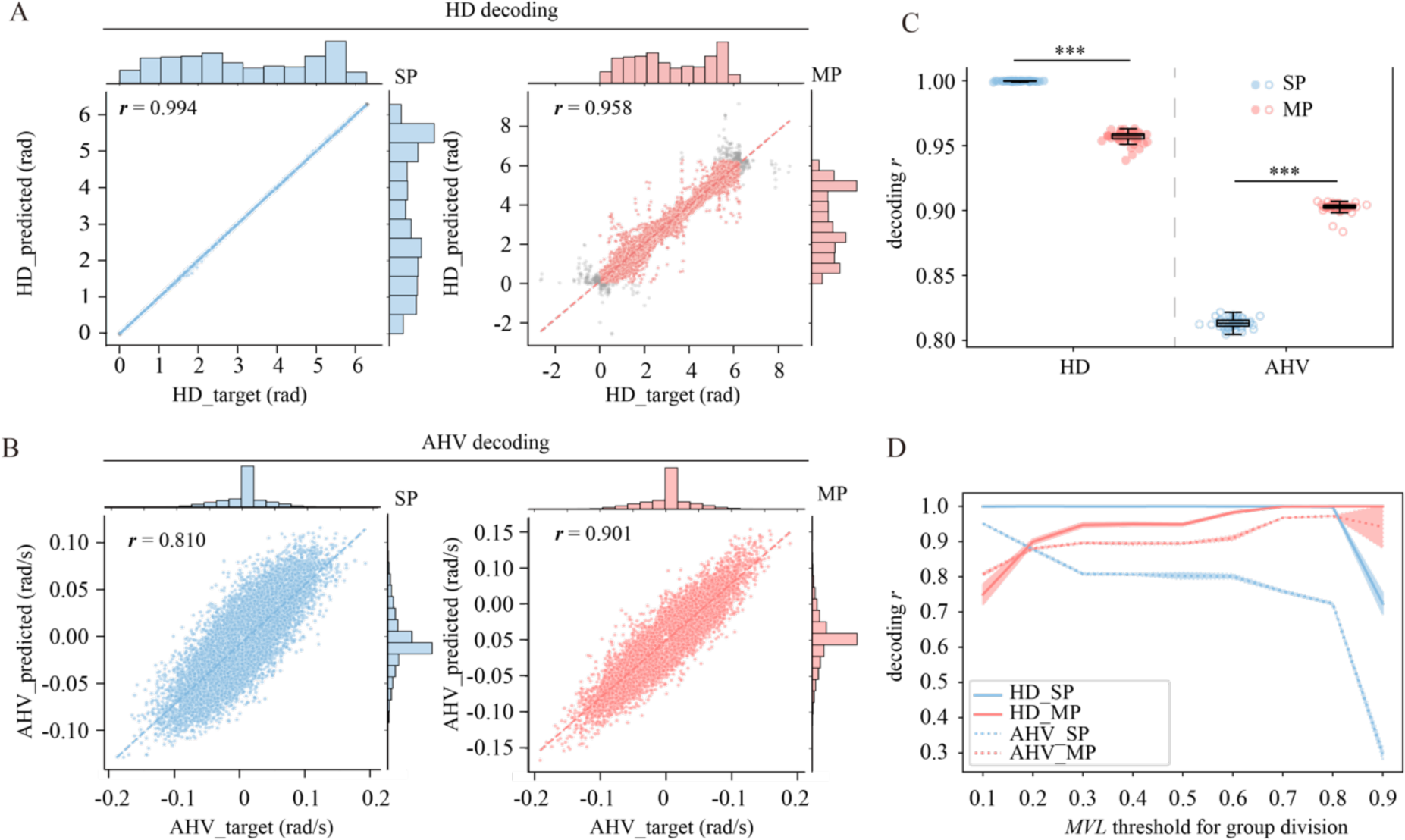
Dissociation of SP and MP unit populations in HD and AHV encoding. (A) – (B) Scatter plots and marginal distributions (histograms) of actual HD (A) and AHV (B) values versus predicted values from ANNs. The *r* values denote Pearson’s correlations between the actual and predicated values. Dashed line denotes the fitted correlation line. Gray dots outside the range of 0∼2π in (A) resulted from correction by shifting 2π when difference between the actual and predicted values exceeded π. (C) Scatter and box plots showing decoding *r* values for unit type (SP versus MP) by interdependent variable (HD versus AHV) across 50 simulated sessions with random seeds. ***: *p* < 0.001. Error bar denotes standard deviation. (D) Changes in decoding *r* values when using different *MVL* thresholds to divide all units into SP (units with *MVL* smaller than the threshold) and MP (units with *MVL* larger than the threshold) populations. Solid and dashed lines represent the mean decoding *r* values for HD and AHV, respectively, and shaded areas represent standard deviation.

In summary, the findings from the computational modeling revealed two distinct populations of units, where units with mixed preferences for various HD slightly compromised their HD encoding capacities to enhance their proficiency in capturing the temporal dynamics of HD, specifically AHV.

### Dissociation between SP and MP neurons in encoding HD and AHV in mice’s HD system

Interestingly, both SP and MP neurons selective to HD have been discovered across various brain regions, including the anterior dorsal nucleus (ADn), Postsubiculum (PoS), retrosplenial cortex, and entorhinal cortex (Hennestad et al., 2021; Taube, 2007), with focus on SP neurons possibly for their majority and simplicity in HD selectivity and ease in computational modeling (Johnson et al., 2005; Wilson, 2023). Inspired by the aforementioned computational modeling, here we delved into the neural representations of HD and AHV in the PoS, the primary cortex of the HD system in mice (Clark and Taube, 2012; Sharp et al., 2001; Winter and Taube, 2014), where neurons attune to both HD and AHV (Sharp, 1996). Utilizing the neural data from CRCNS.org (Peyrache et al., 2015), which includes sorted single-unit firing activities of PoS neurons during mice’s free foraging, we observed that 78% of recorded neurons exhibited significant HD tuning, within which 64% were also responsive to AHV. This higher proportion of neurons conjunctive coding of both HD and AHV compared to previous reports (Sharp, 1996; Taube et al., 1990a) likely stemmed from the inclusion of MP neurons in our study, which were traditionally excluded from analyses. Fig. 4A illustrates neurons with the highest (SP) and lowest (MP) *MI_HD_* from an example recording session, showing their tuning curves for HD similar to the units in the RNN (Supplementary Fig. 1A).

**Figure 4.**
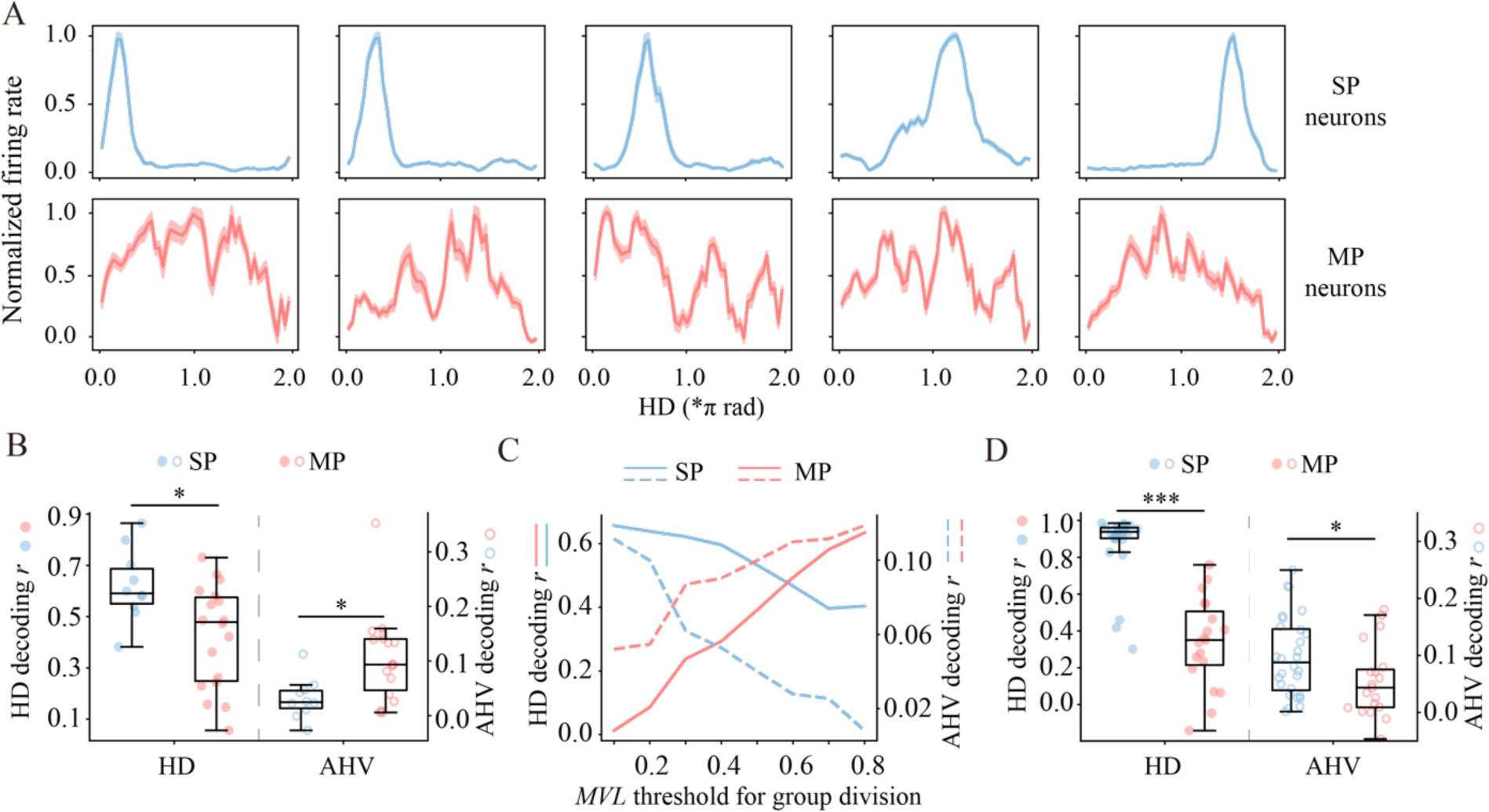
The dissociation of encoding HD and AHV between SP and MP neurons both in the cortex (PoS) and thalamus (And) of mice’s HD system. (A) Top: Tuning curves of five neurons with the highest *MI_HD_*; Bottom: Five neurons with the lowest *MI_HD_* in the PoS from a representative session. (B) Scatter and box plots of SP versus MP neurons in encoding HD versus AHV in the PoS. *: *p* < 0.05; Error bar: standard deviation. (C) Changes in decoding *r* values when using different *MVL* thresholds to divide all neurons from PoS into SP neurons (*MVL* smaller than the threshold) and MP neurons (with *MVL* larger than the threshold) populations. The solid and dashed lines represent the mean decoding *r* for HD and AHV across all sessions, respectively. (D) Scatter and box plots of SP versus MP neurons in the ADn in encoding HD versus AHV. ***: *p* < 0.001; *: *p* < 0.05.

To investigate whether the algorithm discovered in the computational modeling applies to biological systems, we first classified PoS neurons into either SP or MP categories using a two-step classification approach with a strictness parameter α (for details, see Method). At α = 0.1, the categorization aligned closely with visual inspection (Supplementary Fig. 2A and 2C). Employing the same procedure as used in the RNN, we observed a distinct pattern of encoding for HD and AHV between SP and MP neurons (Fig. 4B). Specifically, the SP neurons encoded HD more effectively than the MP neurons (0.62 versus 0.43, *p* < 0.02, Mann-Whitney U Test), whereas for AHV encoding, the trend was reversed (0.03 versus 0.10, *p* < 0.02, Mann-Whitney U Test). This dissociation was further confirmed by a significant two-way interaction of neuron type (SP versus MP neurons) by encoded variable (HD versus AHV) (F(1, 26) = 12.27, *p* < 0.001). In addition, this dissociation was not attributed to neuron classification criteria, as the interaction persisted under stricter (α = 0.05; F(1, 25) = 9.35, *p* < 0.005) or more lenient (α = 0.2; F(1, 27) = 15.34, *p* <0.001) parameters (Supplementary Fig. 2F). Another way of showing the dissociation is to calculate *MVL* values for all neurons and then explore how decoding capacities changed with different *MVL* thresholds for separating neuron populations (Fig. 4C). In line with the pattern observed in the RNN model, a lower *MVL* threshold around 0.2 could led to a superior AHV encoding capacity in MP neuron population. Therefore, the biological systems apparently adopt a similar algorithm for encoding the interdependent variables, with SP neurons specialized in HD encoding and MP neurons excelling in AHV encoding.

A more profound question is whether the functional division of labor in encoding HD and AHV by SP and MP neurons observed extends beyond the PoS and serves as a general principle in the HD system. To address this question, we examined HD and AHV encoding within mice’s ADn, which relays sensory information from peripheral nuclei to the PoS. Employing the same procedure as used in the PoS, we found a similar dissociation pattern in the ADn, as evidenced by a significant two-way interaction between neuron type (SP versus MP neurons) by encoded variable (HD versus AHV) (F(1,45) = 59.77, *p* < 0.001), although the SP neurons outperformed the MP neurons in both HD and AHV encoding (HD: 0.88 versus 0.34, *p* < 0.001, Mann-Whitney U Test; AHV: 0.10 versus 0.06, *p* = 0.04; Mann-Whitney U Test) (Fig. 4D). Taken together, the functional division of labor in encoding HD and AHV between SP and MP neurons might represent a general principle within the HD system, emphasizing the distinct advantage of MP neurons in AHV encoding.

### MP units create high-resolution HD neural manifolds

In both computational modeling and empirical experiments, we revealed a general principle that SP neurons were more adept at encoding HD, whereas MP neurons excelled in encoding AHV. This functional dissociation raises a pertinent question regarding underlying mechanisms that facilitate neurons with MP tuning to exhibit a heightened capacity for AHV encoding compared to their SP counterparts. Besides the difference in mixed selectivity to multiple discontinuous intervals of HD versus pure selectivity to only one HD interval, a prominent distinction lies in the tuning curve width for HD, characterized by the full width at half maximum (FWHM), which provides a concise metric of the neuron’s tuning sharpness for HD. Clearly, MP neurons’ width, indexed by an average of the FWHM across all peaks, was significantly narrower than SP neurons’ width (RNN: 1.29 versus 2.14, *p* < 0.001; PoS: 0.82 versus 1.05, *p* < 0.001; ADn: 0.79 versus 1.28, *p* < 0.001). Recent studies have suggested that the width of a neuron’s tuning curve plays a critical role in shaping the dimensionality of neural representation spaces (De and Chaudhuri, 2023; Kim et al., 2020; Kriegeskorte and Wei, 2021; Langdon et al., 2023), which in turn influences the functionality of neural networks. Thus, here we delved into the impact of tuning curve’s width on HD and AHV encoding. Specifically, we examined all units within the RNN, which allowed us to systematically explore this aspect while mitigating both inherent noise and potential sampling biases in traditional empirical studies.

First, we explored the relationship between units’ width of tuning curves and their capacity of encoding HD and AHV. To eliminate potential confounding biases from mixed selectivity in MP units, here we focused on the variance of width within SP units. We found that the tuning curve width of SP units was positively correlated with *MI_HD_* (r = 0.89, *p* < 0.001), suggesting that units with broader tuning curves more effectively encoded HD (Fig. 5A, blue triangles). In contrast, a negative correlation was observed between the tuning curve width and *r_AHV_* (r = -0.72, *p* < 0.001), suggesting that units with narrower tuning curves were more adept at encoding AHV (Fig. 5A, blue circles). This observation suggests a distinct role for tuning curve width in HD and AHV encoding, aligning with the empirical evidence that MP units, characterized by even narrower tuning curve width than the smallest width in SP units, excelled in AHV encoding (Fig. 5A, red circles).

**Figure 5.**
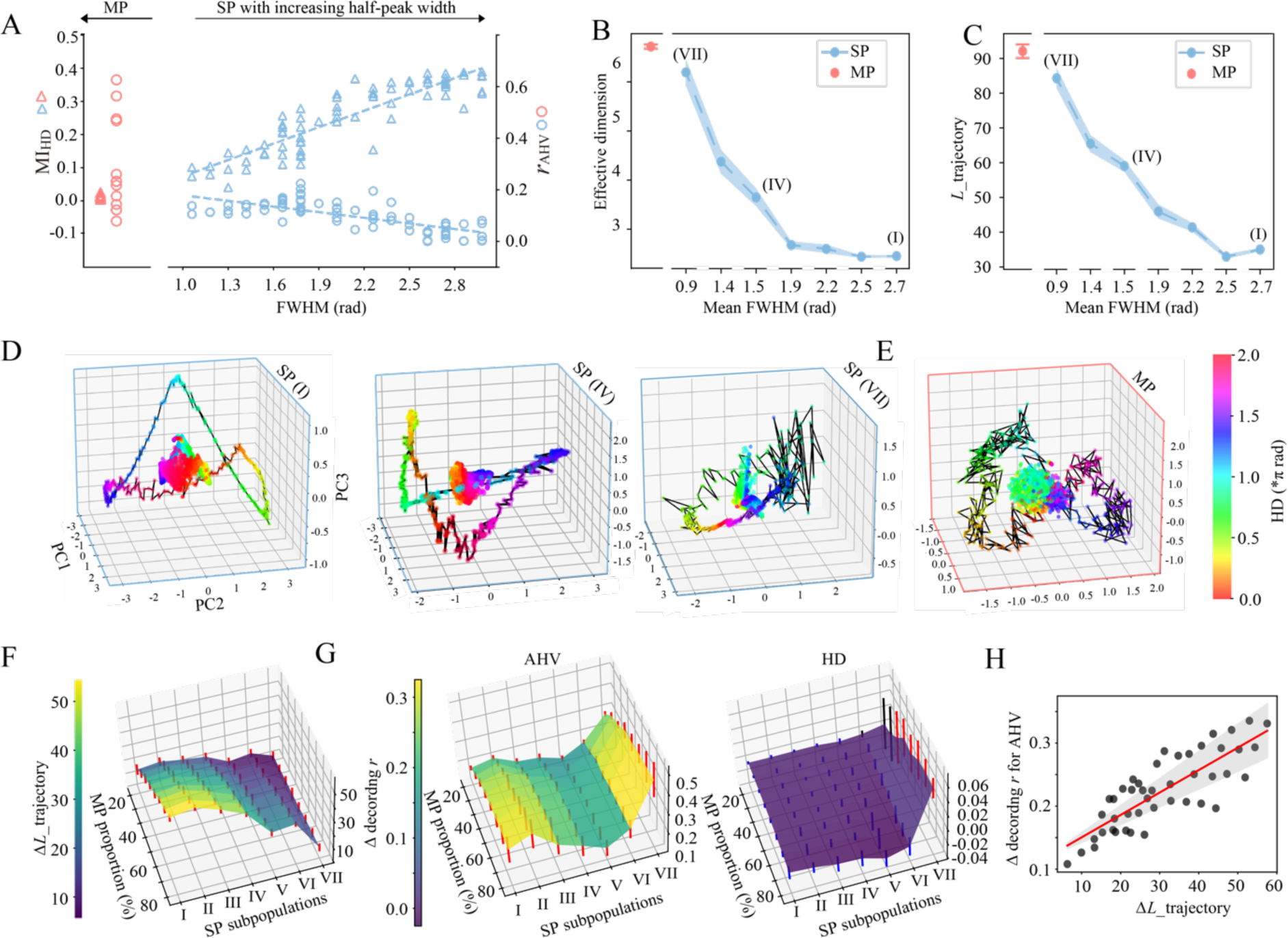
Advantages of MP units for effective dimensionality and high-resolution manifold representation. (A) Relationship between *MI_HD_*, *r_AHV_* and response profiles of RNN units. Blue triangles represent *MI_HD_* and blue circles represent *r_AHV_* for SP units. The dotted lines represent the linear fit between FWHMs and *MI_HD_*/ *r_AHV_*. Red triangles and circles represent *MI_HD_* and *r_AHV_* for all MP units, respectively. (B) Effective dimensions for 7 SP subpopulations and 1 MP subpopulation across 20 simulated sessions with different random seeds. X-axis denotes mean FWHM for each SP subpopulation. (C) Trajectory length (*L*_trajectory) of the HD manifold in the original 11-D space for 7 SP subpopulations and 1 MP subpopulation across 20 simulated sessions with different random seeds when dividing 0∼2π into 360 bins. (D) – (E) HD manifolds for SP subpopulation I, IV and VII as depicted in (B) and (C), and for MP subpopulation, embedded in the representational space defined by the first three principal components of their own populational activity. Lines connecting discrete colored dots represent the HD manifold obtained by averaging each unit’s firing rates within each 1° HD bin. Collapsed point clouds are the projections of neural states corresponding to all timesteps in a simulated session. These projections are normalized by the maximum embedding projections across all three component axes to distinguish them from the mean HD manifold. The color of the dots and points indicates the value of HD, as shown in the color bar in (E). (F) – (G) The change in trajectory length (△*L*_trajectory), decoding *r* for AHV and HD, for a representative simulated session when randomly replacing different proportions of SP units with MP units for each SP subpopulation. Error bars represent standard deviation over 20 trials of random replacement. Red and blue error bars represent mean values significantly larger and smaller than 0, respectively, while black error bars indicate no significance compared to 0. (H) Relationship between the change in HD manifold trajectory length and the change in decoding *r* for AHV while randomly replacing SP units with MP units for SP subpopulation I to VI.

To illustrate the mechanistic effect of tuning curve width, we examined the characteristics of the neural representation space constructed by SP units with varying tuning widths. Specifically, we sorted all SP units in the RNN based on their FWHMs, and then evenly classified them into 7 subpopulations, each of which contained 11 units. We assessed the effective dimensionality (Del Giudice, 2021; Farrell et al., 2022) of the neural representation space constructed by each subpopulation, with higher dimensionality implying a more diverse and potentially more nuanced representation encoded by the neuron population. As expected, the effective dimensionality exhibited a monotonic increase with the narrowing of the units’ tuning curve widths (F(6, 13) = 1485.4, *p* < 0.001, Fig. 5B), suggesting that units with narrower tuning curves contributed to a neural representation space of a higher dimensionality. To visualize the HD manifold embedded in the neural representation space for each subpopulation, we first z-scored the firing rate for all units to eliminate bias introduced by differences in response strength across units. Then, we averaged firing rates for each unit within each HD bin of 1° to obtain mean HD manifold, which were further projected onto a 3-D neural subspace defined by the first three principal components from the same subpopulation (see Methods). Fig. 5D shows the HD manifolds depicted by colored dots threaded by black lines, which formed a 1-D ring representing HDs from 0 to 2π. Critically, the HD manifolds generated by unit subpopulations with varying tuning curve widths significantly differed in degree of smoothness (i.e., local structure). The HD manifold by units with narrower width appeared jagged, with abundant twists and turns (Fig. 5D, Right), indicative of higher dimensionality. In contrast, the HD manifold by units with broader tuning curves was smoother (Fig. 5D, Left), indicating lower dimensionality. Because AHV is the temporal derivation of HD, the rotation speed of AHV equates to the change in HDs over an extremely short period. In terms of neural geometry, the proximity between two successive HDs signifies AHV. Therefore, greater local structure complexity corresponds to a longer distance between two successive HDs, hereby providing more accurate representation of AHV. To quantify this intuition, we measured the trajectory length along the HD manifold for each of the aforementioned 7 SP subpopulations by summing the Euclidean distances in the original 11-D space between successive HDs at 1° intervals (see Methods). We found that manifold trajectory length increased as tuning curve widths narrowed (F(6, 13) = 2138, *p* < 0.001) (Fig. 5C), suggesting that units with narrower tuning curves provide finer resolution for AHV representation. Accordingly, the most effective strategy to narrow the width of tuning curves is to introduce MP units, which possess tuning curves embedding peaks with very narrow widths. Not surprisingly, the neural representation space constructed by MP units exhibited the highest effective dimensionality (Fig. 5B, red), the longest distances between neighbor HDs (Fig. 5C, red), embedding the HD manifold with highest local structure complexity (Fig. 5E) and hereby achieving the highest resolution in representing AHV. Interestingly, when randomly replacing 20%∼80% of the SP subpopulation with MP units (see Methods for details), a consistent increase in manifold trajectory length was observed (Fig. 5F). For most of the SP subpopulations (from I to VI), this increase was also significantly correlated with increased values of the decoding *r* for AHV (r = 0.84, *p* <0.001) (Fig. 5H). Conversely, the decrease in the decoding *r* for HD after being replaced with MP units was almost negligible (mean change in decoding r: -0.008 for HD versus 0.22 for AHV) (Fig. 5G, right). Therefore, these results collectively demonstrated that MP units effectively expand the dimensionality of neural representation spaces and increase local richness of HD manifold, facilitating high-precision representations of AHV, the temporal derivation of HD.

## DISCUSSION

To explore the simultaneous encoding of the interdependent features of HD and AHV in the brain, we developed a novel methodological approach. Specifically, we utilized a computation model of HD systems (Cueva et al., 2019) for exploratory analysis, followed by empirical validations to confirm the biological plausibility of the algorithm identified in the artificial HD system. Specifically, the model only simulated the core functionality of the HD system that leverages current HD and AHV signals to predict upcoming HD values, hereby minimizing confounding factors often encountered in biological systems. In addition, unlike traditional HD models that typically predefine SP units in a circular arrangement (Clark and Taube, 2012; Skaggs et al., 1995; Vafidis et al., 2022; Zhang, 1996), this model is hypothesis-neutral, allowing for exploring emergent algorithms. Through this methodology, we identified two distinct neuron populations: MP neurons, which compromised their HD encoding capacity to better capture AHV dynamics, and SP neurons, which predominantly encoded HD. Further analyses on the neural geometry constructed by MP and SP neurons revealed that tuning curve widths played a critical role in determining the local complexity of the neural manifold, which in turn affected the dimensionality of the neural representation space to modulate the representation precision based on task demands. Given the discovery of the conjunctive coding of interdependent features in domains other than navigation, such as neurons in the primary motor cortex that encode both hand position and velocity (Paninski et al., 2004; Wang et al., 2007), our study offers a new framework for examining the representation of interdependent features across various neural domains.

Our finding of the functional dissociation in the encoding of HD and AHV by SP and MP neurons complements recent computational modeling of the granular retrosplenial cortex that focused exclusively on SP neurons (Brennan et al., 2021). Our study advanced this finding by identifying MP neurons, which are distinct from SP neurons not only in their mixed selectivity to HDs but also in the preference for AHV encoding over HD. Further neural geometric analysis revealed that the multiphasic characteristic of MP neurons contributed to the formation of locally jagged and complex neural manifolds, leading to higher effective dimensionality. This complexity enables a longer trajectory length that more effectively captured the temporal dynamics of AHV. A similar association between local richness of manifolds and dimensional elevation has been observed in simulated population activity of the visual cortex in encoding parameterized one-dimensional stimulus, where a slower eigen-spectrum decay in neural codes implied additional dimensions for encoding fine stimulus details (Stringer et al., 2019). Such detailed representations permit a more precise encoding of changes in HDs over short periods of time, thus providing a foundation for the superior precision in AHV encoding.

The key factor for expanding the dimensionality of neural representational spaces is the width of tuning curves. Research has repeatedly shown that, for one-dimensional features, narrower tuning curves not only enable individual neurons to encode a greater volume of information (Zhang and Sejnowski, 1999) but also contribute to increasing the linear dimensions of neural populations (De and Chaudhuri, 2023; Kriegeskorte and Wei, 2021) and producing a more reliable signals for low latency communication (Lenninger et al., 2023). Note that this characteristic has previously been discussed primarily in the context of SP neurons, which exhibit a great variance in their tuning curve widths. In our study, MP neurons displayed an average tuning width that closely aligned with that of the narrowest SP neurons, and constructed a neural representational space with higher dimensionality. Hence, the differentiation in AHV representation by SP and MP neurons may not be qualitive in nature; rather, MP neurons may simply provide a more efficient representation strategy. This leads to the hypothesis that MP neurons could have evolved from SP neurons by modifying the response profiles of SP neurons from a single HD focus to encompass multiple HDs. This transformation in response profiles may provide an example, marking a shift from neurons with pure selectivity to those with mixed selectivity.

Because HD and AHV are interdependent, with AHV providing essential information for updating HD, the encoding challenge for these two interdependent features lies in balancing the preservation of specificity for individual features while maintaining their interdependency nature. This challenge thus represents a significant departure from the encoding of independent features. Specifically, to preserve specificity, the HD system adopted a sparse-like coding scheme, where SP neurons specialize in HD encoding and MP neurons excel in AHV encoding. This sparse-like coding scheme may serve as an efficient strategy to keep individual features distinct and minimize mutual interference, thereby maintaining the balance between resistance to inter-neuron interference and the ability to generalize (Beyeler et al., 2019). Given that HD and AHV information may be utilized differently at subsequent neural processing stages, the sparse-like coding strategy also capitalizes on its energy efficiency and simplifies the access for readouts (Olshausen and Field, 2004; Takiyama, 2015). In this context, the behavior of SP and MP neurons resembles that of pure selectivity neurons.

In contrast, to maintain their interdependent relation, the HD system also employed a dense-like coding scheme, with both SP and MP neurons responding to HD and AHV. Furthermore, although MP neurons exhibited the narrowest tuning curve width, the difference in width between MP and SP neurons was not qualitatively significant. In other words, without MP neurons, SP neurons alone could competently encode AHV, albeit not as effectively as when a small fraction of MP neurons was introduced. In this context, SP and MP neurons do not differ qualitatively; rather, they operate similarly to neurons with mixed selectivity. Recent studies (Kira et al., 2023; Ledergerber et al., 2021; Rigotti et al., 2013) have shown that mixed selectivity plays a key role in constructing higher-dimensional representational spaces (Kriegeskorte and Wei, 2021; Ma et al., 2023; Rigotti et al., 2013; Sussillo and Abbott, 2009), potentially facilitating the integration of interdependent features into a cohesive whole. Therefore, the coding scheme for interdependent features appears to be a hybrid of both dense and sparse coding schemes, aiming at achieving the balance between specificity and interdependency, marking a significant departure from the encoding of independent features and thus advocating future research to delve into this emerging field.

## Acknowledgements

We deeply appreciate Dr. Wei and Dr. Cueva for sharing their code of the RNN model on the HD system and Dr. Peyrache for sharing their neurophysiological data. This work was supported by Beijing Municipal Science & Technology Commission & Administrative Commission of Zhongguancun Science Park (Z221100002722012), and Double First-Class initiative Funds for Discipline Construction of Tsinghua University (J.L.).

## Author Contributions

J.L. conceived the study and provided advice. D.C. analyzed RNN data. D.C. and T.L. analyzed the mice data. T.L. designed the ANN for population decoding. D.C. and J.L. wrote the manuscript.

## Data and Code availability

The data that support the findings of the present study are available at CRCNS.org [https://portal.nersc.gov/project/crcns/download/th-1]. All analyses reported in the present study were conducted with customized code written in Python (version 3.9) and Matlab (version 2021a). The code will be available from the corresponding author (J.L.) upon request.

## Materials and METHODS

### Recurrent neural network

The recurrent neural network (RNN) for integrating angular velocity into head direction was trained using the procedure described in Cueva et al., 2019. As illustrated in Fig. 1A, the RNN contained N fully recurrently connected units (*N* =100 or *N* = 64 in this study) with connection weights 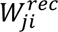. Each unit also received 3 external inputs through weight matrix 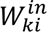 and projected outputs through weight matrix 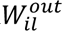 to 2 output units. The 3 inputs were cos θ*_t_*, sin θ_&_ and angular velocity *AHV_t_*, which were divided into two phases:

1. In the first phase lasting 10 timesteps, *AHV*_&_ was 0, cos θ_1_ and sin θ_1_ were determined by initialized HD θ_0_.
2. In the second phase for all subsequent timesteps, cos θ_1_ and sin θ_1_ were 0, and *AHV*_&_ at each timestep was simulated as *AHV_t_* = σ*X* + *momentum* ∗ *AHV_t_*_23_ with σ = 0.03 and *momentum* = 0.8. X is a zero mean Gaussian random variable with standard deviation of one. The seeds of the random variables changed in different simulations.

The outputs were the updated cos θ_1_ and sin θ_1_ of the current step, where θ_1_ was the exact integral of all previous angular velocity.

The dynamics of each recurrent unit in the RNN was updated using a biologically-plausible firing rate model, which assumes that firing rate responds instantaneously to synaptic input, but synaptic conductance change slowly as shown by the following simplified input-output characteristics and differential equation for synaptic input:

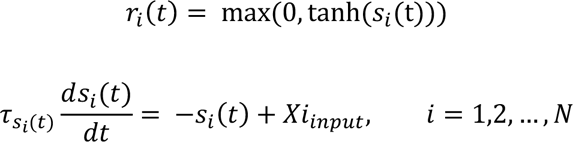

Where τ_*si*(*t*)_ was set to be 250ms for all unit and *Xi_input_* is the summation from recurrent inputs, external inputs and constant bias:

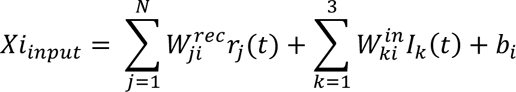

The activities of two output units were given by:

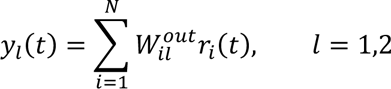

The model was trained with the hessian-free algorithm under the *L*_2_ regularization on firing rates of both recurrent and output units to minimize the mean-squared error between the target and the theoretical head direction:

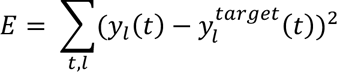

In practice, the network activities were simulated using Euler method for 500 timesteps of 25ms duration. Parameters were optimized by minimizing mean-squared error across 500 trials in parallel to increase training efficiency. Half of the trails were allowed to have duration of *AHV* = 0 for up to one-third of the trial length. The model was trained for 500 epochs in total, and model parameters for epoch 500 were saved to generate dataset for further analyses.

## DATASET

This study analyzed datasets of firing activities from both artificial and biological neural network. The first dataset consisted of firing activity from recurrently connected units in an RNN model for specific training epoch during 200 new trials with 1000 timesteps per trial (i.e., 200000 samples in total for each unit with a sampling interval of 25ms). All results were derived from the firing activities generated by RNN model with parameters from training epoch 500. The second dataset contained recorded neural spiking activity from mouse brains using silicon probes, shared by Peyrache et al. (2015) and obtained from the CRCNS website. Neuron region labels (ADn and PoS) were extracted for each neuron from detailed session information provided with the dataset. Thirty-seven complete recording sessions across 6 mice were included for further analysis. For analysis of HD and AHV decoding from the neuronal populations in ADn and PoS, only sessions with >30,000 samples (ensuring small standard deviations across cross-validation splits, Supplementary Fig. 2E) and containing at least 3 neurons in both the SP and MP groups were used.

### Preprocessing of behavioral and neural data in mice

The original head direction angle lists during awake states were first converted into accumulated angles from a circular distribution. To reduce errors caused by video interlacing, the accumulated angles were smoothed by a Gaussian kernel with σ = 200ms and moving step size = 25ms. In practice, a sliding Gaussian window *G*(*t*) of width 6 σ was used to increase processing speed while retaining sufficient resolution for further analyses. Angular velocity lists were obtained by convolving the accumulated angle lists with a Gaussian derivative sliding window defined as *G*^?^(*t*) = *G*(*t*) ∗ (*x* − μ)/σ^2^ for every 25ms bin, followed by normalizing. This convolution of the accumulated angles with the Gaussian derivative window enabled efficient and accurate computation of the angular velocities.

To align the timelines between the neural firing and behavioral data, the original timestamps for spikes of each sorted neuron were convolved with the same Gaussian sliding window *G*(*t*) as above, with width 6 σ and moving step size = 25ms, followed by summation.

### Analysis of HD and AHV units/neurons

We utilized the same method to firstly construct HD and AHV tuning curves both for units from the RNN model and neurons from mice recordings. The 0 to 2π angular radian range was divided into 50 equal bins. For each neuron, the HD tuning curve was obtained by calculating the mean firing rate for each angular bin after averaging the instantaneous firing across all timepoints where the animal’s HD direction fell within that bin. For AHV tuning curves, mean firing rates for bins of 0.2 rad/s were derived by averaging instantaneous firing for timepoints when the AHV was within each bin. Tuning curve standard deviations were determined through 50 permutations, constructing the tuning curves each time using a randomly selected half of the full sample points. We excluded twelve units in RNN from further analysis (Supplementary Fig. 1A, units with gray tuning curves) due to low tuning stability or extremely low activities (mean firing < 0.0001Hz, Supplementary Fig. 1C) throughout the simulation process.

The strength of HD tuning for each neuron was assessed by computing the mutual information between HD and the averaged firing rates across different HD bins defined as (Skaggs et al., 1993):

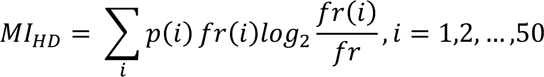

where *fr*(*i*) and *p*(*i*) are the mean firing rate and the probability that the animal’s head pointed in the th head direction bin, and *fr* is the overall mean firing rate of the unit/neuron.

The strength of AHV tuning were measured as *r_AHV_*, defined as the maximum of *r_AHV _pos_* and *r_AH _neg_*, where *r_AHV _pos_* and *r_AHV _neg_* are the absolute values of the Pearson’s correlation coefficient between firing rate and angular head velocity when velocity is greater than zero or less than zero, respectively. The significance of the tuning for both HD and AHV were determined through a shuffling process. The chance-level distribution of *MI_HD_* and *r_AHV_* were derived from 1000 repetitions with shuffled condition. For each repetition, the firing rate time series was shifted by a random interval of at least 30s, with the end of the trial wrapped to the beginning. Significantly tuned HD and AHV units and neurons were defined when calculated *MI_HD_* and *r_AHV_* values were higher than 99th percentile of the shuffled distribution.

### Classification of SP and MP HD units/neurons

In the RNN model, SP and NO HD units were segregated automatically by two phases of linear fitting between *MI_HD_* and *r_AHV_* as shown in Fig. 1C or Supplementary Fig. 2D. The linear regression line was obtained by least squares error fitting within each phase separately.

In mice recordings, a two-step classification approach was used to categorize SP and MP HD neurons. A neuron was defined as SP HD if it met the following two criteria:

1. Its HD tuning curve contained no bins within the range of interest that showed a mean firing rate larger than half of the maximum response across all HDs. The range of interest was defined as the complementary set to the range spanning the half-peak bins detected on either side of the peak bin.
2. Its HD tuning curve had fewer than α of bins in the range of interest with mean firing rates exceeding 20% of the maximum response. The range of interest here was the complementary set to the range spanning the 20% peak bins detected on either side. Three values of α were tested in this study, including α = 20%, 10%, 5% corresponding to increasingly strict criteria for categorizing a neuron as SP.

In order to quantify the multi-peak nature of the tuning, we calculated mean vector length (*MVL*) for each unit/neuron, which typically was used to describe the strength of HD tuning for conventional SP neurons. *MVL* was defined as following:

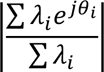

Where *j* is the imaginary unit, θ_(_ is the head direction angle in radians associated with the *i*-th bin, and λ_*i*_ is the mean firing rate in that bin. By applying *MVL*, a metric normally used for SP tuning, to MP neurons, we could quantify the degree of multiphasic in their tuning curves. The lower the *MVL* value, the more uniform distributed in amplitudes for the tuning curve.

### Feedforward neural network

We assessed the encoding capability of neuronal populations using a five-fold cross-validation with well-trained artificial neural networks (ANNs). The underlying hypothesis was that a feedforward ANN, known as a universal approximator, could extract the maximal information from the population activity patterns.

Two ANNs were trained for each population group (SP or MP) in both the RNN and mice datasets - one for predicting HD and one for predicting AHV. The input to the ANNs was the instantaneous firing rates of the neuronal populations. The output was either the corresponding HD or AHV value at each timepoint.

Both ANNs had 3 fully connected hidden layers with 64, 128, and 64 units respectively. Each unit in a layer was computed from a linear combination of the previous layer’s outputs passed through a rectified linear activation function called ReLU (Glorot et al., 2011). The Adam optimization algorithm was used (Kingma and Ba, 2015), with weight decay set to 5e-2 and 5e-3 for the HD and AHV ANNs, respectively.

### Measurement of FWHM for SP tuning and MP tuning curves

To estimate the full width at half maximum (FWHM) as the main feature of the response profile, all tuning curves were firstly normalized by subtracting their minimum value and then underwent further processing. For SP tuning curves, we first located the peak of the tuning curve along with its corresponding angle value. From the peak, we then proceeded to find the first angle values on both the left and right sides where the tuning value was less than half of the maximum value. These angles were denoted as HD_HM_left and HD_HM_right. The FWHM was then measured as the angular difference between HD_HM_left and HD_HM_right (in radians). For MP tuning curves, local maxima were first detected using find_peaks function, with a distance parameter set to 10 and 8 for MP units/neurons from the RNN model and mice data, respectively. Then local valleys were detected on both sides of each local maximum. Qualified peaks were defined when at least one side of the peak had valley values lower than half of the peak value. Subsequently, similar to procedure for SP tuning curves, HD_HM_left or/and HD_HM_right was determined for each qualified peak when applicable. The FWHM of each qualified local peak was calculated as the angular difference between HD_HM_left and HD_HM_right when half maximum was found on both sides of the peak. If only one side of the peak had a half maximum, the FWHM was calculated as twice the angular difference between HD_HM_left/ HD_HM_right and the angle corresponding to the local peak. Finally, the mean of all qualified FWHMs was calculated as the FWHM of the MP unit/neuron. On average, 2.2, 3.5 and 3.3 qualified peaks were found for each MP unit/neuron in the RNN model, ADn and PoS, respectively.

### Effective dimensionality

According to (Farrell et al., 2022), the effective dimensionality (ED) of a set of S points *V* = [*V*_*sk*_] ∈ *R*^*SXd*^ with embedding dimension *d* is

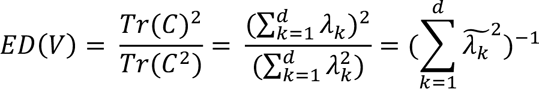

where *Tr* is the trace operation for matrices, *C* is the covariance matrix of *V* and 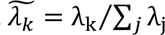 are the normalized eigenvalues of *C*. Therefore, to calculate ED for each subpopulation of 11 units analyzed in the RNN model, the first step was to construct an activity matrix *T* of size 11*200000, where each row represented the activity time series of a unit in that group during a simulation session. Then, PCA analysis was performed on the transpose of the *T* to obtain explained variance ratio of each PCA component, i.e., 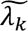. Thus, the ED could be calculated according to the above equation.

### Characterization of the HD manifold in high-dimensional activity space of RNN

To eliminate potential artifacts introduced by the metabolic cost used in training the RNN and to concentrate solely on how response profiles could affect the high-dimensional geometry of the unit population, we applied z-scoring to the firing rate of each unit before characterizing the HD manifold. Subsequently, we divided the range of 0∼2π into 360 bins and calculated averaged firing rate within each bin for each unit in the population to derive the 11-D mean HD manifold. Finally, trajectory length (*L*_*_trajectroy_*) along the HD manifold was obtained by summing the Euclidean distances between neural states corresponding to each pair of adjacent HD bins. To visualize the geometric characteristics of the high-dimensional HD manifold, we first computed the first three principal components (PCs) for the 11-D neural states corresponding to all simulated timesteps. Then, mean HD manifold embedded in this lower 3-D neural representation space was obtained by projecting the original 11-D neural states onto the three PC axes, as shown in Fig. 5D and E. Additionally, down-sampled neural states were projected in the same three PCs coordinate system but were normalized by dividing the maximum of the projected values of all three PC axes to disambiguate the visualization of the mean HD manifold.

To assess how the incorporation of MP units influences the local richness of the HD manifold and the encoding capability HD/AHV for prementioned 7 SP subpopulations with increasing mean FWHM. We randomly replaced *M* (*M* = 2∼9) SP units with that randomly selected from MP subpopulation, resulting in MP units accounting for about 20%∼80% of the subpopulation. Then, we computed the difference in trajectory length (△ *L*_*_trajectory_*), ANNs based decoding *r* for HD and AHV between the mixed group (comprising *M* MP cells and 11-*M* SP cells) and the pure group (comprising 11 SP units of the original subpopulation). This process was repeated 20 times to obtain mean and standard deviation values, as depicted in Fig. 5F and G.

### Software and Statistical analysis

The RNN model training and data generation were performed in MATLAB 2021a (Mathworks), using code generously shared by Cueva et al. (2019). Subsequent analyses and statistical test of the simulated RNN unit activities and recorded neuronal firing were all conducted in Python using the VScode environment.

## Supplementary Materials

**Supplementary figure 1.**
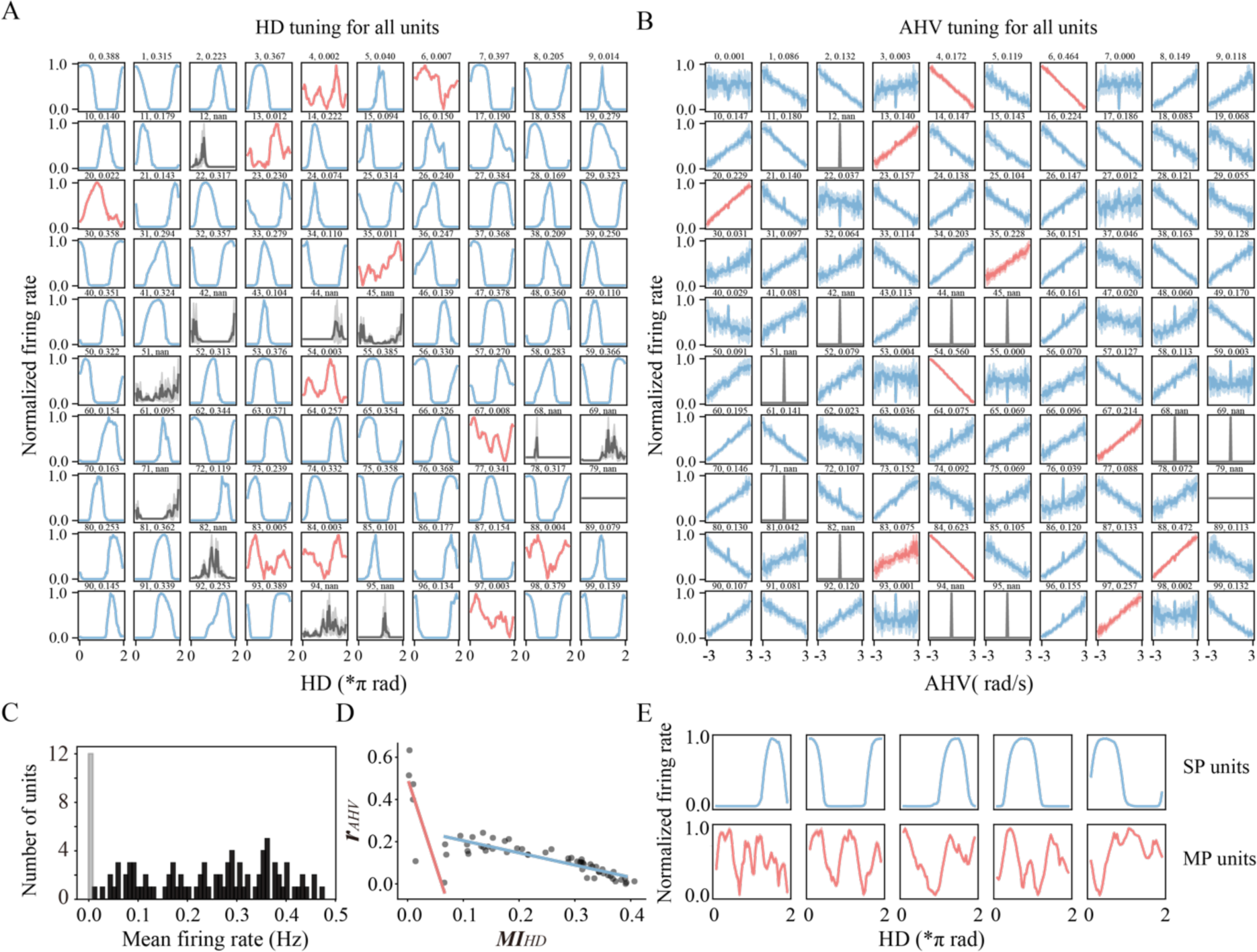
Emergence of single-peaked (SP) and multiple-peaked (MP) units of HD in trained RNNs with network size of both 100 and 64, related to Figure 2. (A and B) Tuning curves of HD and AHV for all units in 100-unit RNN. Units with gray tuning curves are excluded from further analysis due to low mean firing rate and unstable tuning. SP units are represented in blue, while MP units are denoted in red. This color code for SP and MP units remains consistent throughout all subsequent figures. Shaded areas represent the standard deviation of tuning curves obtained from 50 permutations of the original firing and HD sequences. Shaded areas are not visible for SP and MP units due to negligible standard deviation across permutations. Unit number and *MI_HD_* are labeled at the top of each HD tuning plot in (A); unit number and *r_AHV_* are labeled at the top of each AHV tuning plot in (B). In (B), only the tuning curves with AHV values in the range of [-3,3] are shown for a better comparison across units. *MI_HD_* and *r_AHV_* values of *nan* represent units excluded for analysis due to excessively low mean firing rates observed throughout the simulation. (C) Distribution of mean firing rates for all units, gray units in (A and B) are included in the histogram around 0. (D and E) Emergence of two populations in the 64-unit RNN. (D) Relationship between the representations of HD and AHV for 59 units significantly tuned to HD in the 64-unit RNN. Blue linear regression line, *ρ*= -2.995, *p* = 0.064; red linear regression line, *ρ*= -0.254, *P* = 1.12*e-13. (E) Tuning curves of HD for five units with the highest *MI_HD_* (upper row) and five units with the lowest *MI_HD_* (bottom row) in the 64-unit RNN.

**Supplementary Fig 2.**
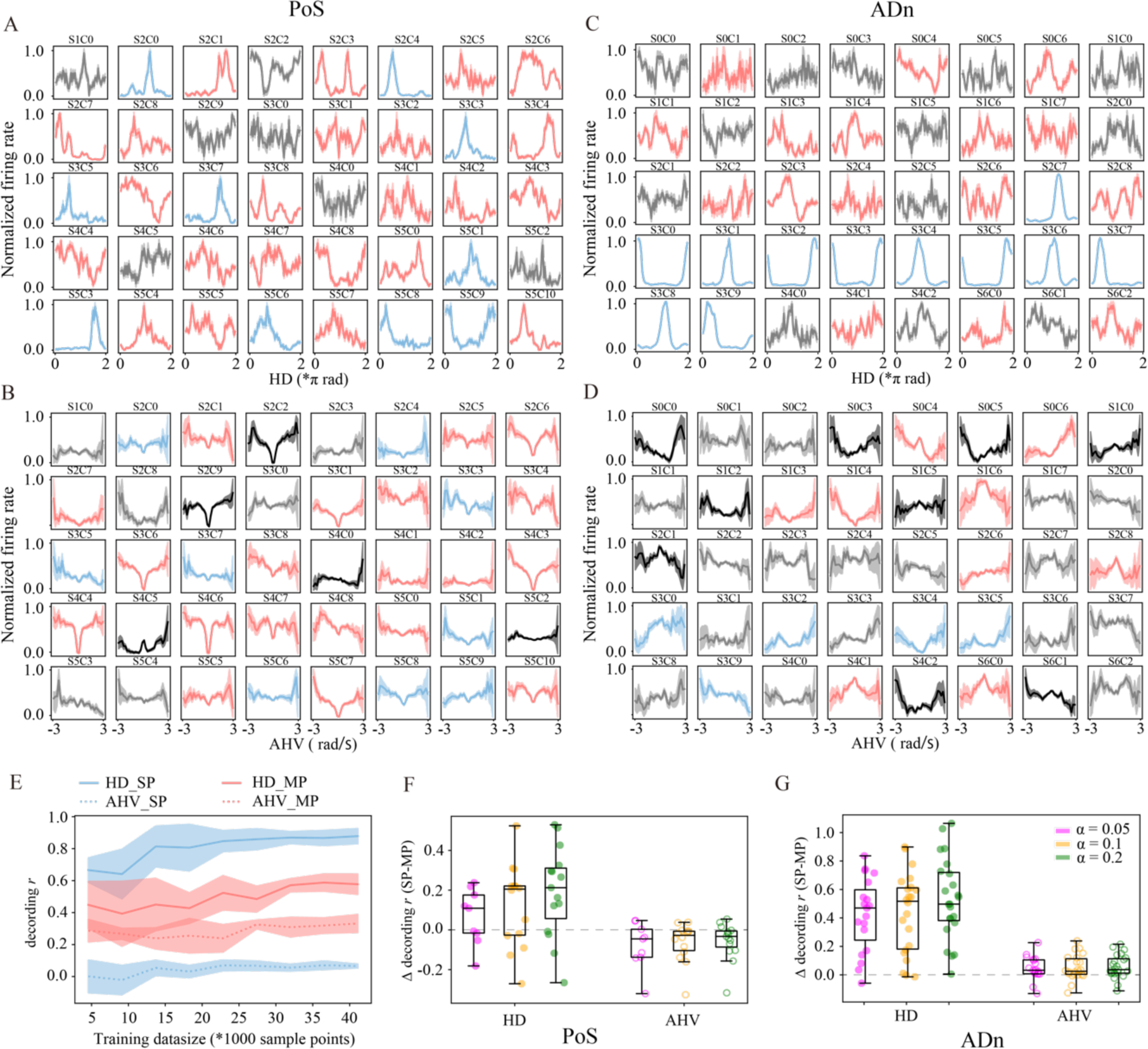
Detailed examples and additional analysis, related to Figure 4. (A and B) HD and AHV tuning curves for neurons recorded in PoS during exemplar session Mouse28-140312. Blue and red neurons in (A) are HD-tuned and classified as SP and MP, respectively. In (B), black, blue, and red neurons are AHV-tuned; among them, only blue and red neurons are also significantly HD-tuned, representing SP and MP HD neurons, respectively. Shaded areas represent standard deviation of tuning curves obtained from 50 permutations of the original firing rates, HD and AHV sequences. Unit shank and cluster are labeled at the top of each tuning plot. (C and D) same as (A and B) but for neurons in ADn during exemplar session Mouse12-120808. (E) Change in decoding correlations when increasing training data size for the illustrated session in (A). The solid and dashed lines represent the mean decoding correlations for HD and AHV, respectively. Shaded areas represent the standard deviation of all five-fold leave-one-out tests. (F) Scatter plot and box plot of difference in decoding *r* between SP and MP neuron populations for HD and AHV across all recording sessions from PoS, with the use of three different strictness parameters α. (G) same as (F) but for recording sessions from ADn. No significant main factor effects or interaction effects were observed for α with three-way ANOVA analysis in both (F) and (G).

